# Defective photosynthetic adaptation mechanism in winter restricts the introduction of overwintering plant to high latitudes

**DOI:** 10.1101/613117

**Authors:** Yue-Nan Li, Yu-Ting Li, Alexander G. Ivanov, Wan-Li Jiang, Xing-Kai Che, Ying Liang, Zi-Shan Zhang, Shi-Jie Zhao, Hui-Yuan Gao

## Abstract

Because of the need for agriculture and landscaping, many overwintering evergreen and biennial species that maintain green leaves over winter were introduced to higher latitudes. The green leaves of introduced overwintering species have to withstand a harsher winter, especially lower temperature, than in their native region of origin. Although the responses and adaptability of photosynthetic apparatus to winter conditions in native overwintering species were widely studied, the experimental results on the introduced overwintering species are very limited. Here, the photosynthetic adaptability during winter was analyzed in two native overwintering species, pine (woody plants), winter wheat (herb), and two introduced overwintering species, bamboo (woody plants), lilyturf (herb). The native species exhibited higher capacity for photosynthetic CO_2_ fixation and lower susceptibility for photoinhibition than introduced species during winter. Photosynthesis related proteins, such as PsbA, PsaA, Rubisco and Lhcb1, were marginally affected in native species, but significantly degraded in introduced species during winter. More interestingly, the PSII photoinhibition was mainly caused by up-regulation of photoprotection mechanism, non-photochemical quenching, in native species, but by photodamage in introduced species. This study indicates that the growth and survival of introduced overwintering species is limited by their photosynthetic adaptability to the harsher winter conditions at high latitudes.

## Introduction

Plants can be divided into deciduous and evergreen (overwintering) species according to their leaves habit during winter. The overwintering species maintain green leaves (or needles) over the winter. To meet the needs of agriculture and landscaping, many overwintering evergreen and biennial species, including trees and crops, were introduced to higher latitudes. The introduction of overwintering species to higher latitudes is more difficult than deciduous species due to the environmental challenges that green leaves of introduced overwintering species has to withstand harsher winter conditions compared to their native region of origin. The survival and avoiding serious injuries of evergreen leaves during winter is critical for the success of introduced species to higher latitudes. In winter, the green leaves of overwintering species have to withstand and cope not only the dehydration and frost damage but also the possible photodamage of photosynthetic apparatus caused by excess light absorption by the chlorophylls in evergreen leaves (Öquist and Huner, 2003). The low temperatures in winter imposes thermodynamic restrictions and slows down the activities of Calvin cycle related enzymes, so not 100% of the light absorbed by the light harvesting complexes (LHC) can be utilized for CO_2_ fixation. The imbalance between the capacity for harvesting light energy and the capacity to dissipate this energy through metabolic activity such as CO_2_ assimilation, can potentially result in generation of reactive oxygen species (ROS) leading to photoinhibition and photooxidative damage of photosystem II (PSII) (Aro *et al*.,1993; Takahashi and Badger, 2011). In addition to PSII, various environmental stress conditions can also cause photoinhibitory damage on photosystem I (PSI) (Sonoike and Terashima, 1994; Sonoike, 1995; Ivanov *et al*., 1998).

The effects of low temperature and high light during winter on photosynthetic apparatus of native overwintering species and their photosynthetic adaptability during winter have been extensively studied (Adams *et al*., 2001; 2004; Öquist and Huner, 2003; Verhoeven, 2014; Míguez *et al*., 2015).

The photosynthetic CO_2_ fixation was almost completely inhibited during winter in evergreen trees (Öquist and Huner, 2003; Russell *et al*., 2009; Savitch *et al*., 2010). These changes were accompanied by significant alterations in chloroplast ultrastructure resulting in a substantial loss of thylakoid grana during winter (Yokono *et al*., 2008; Maslova *et al*., 2009; Silva-Cancino *et al*., 2012). Both PSI and PSI photochemical activities decreased (Ottander *et al*., 1995; Ivanov *et al*., 2001; 2002; Ensminger *et al*., 2004; Robakowski *et al*., 2005) and this was attributed to degradation of a number of PSII and PSI related proteins during winter (Ottander *et al*., 1995; Ebbert *et al*., 2005; Verhoeven *et al*., 2009; Míguez *et al*., 2017). However, the PSII photoinhibition was more pronounced than that of PSI during winter (Ivanov *et al*., 2001; 2006).

The sustained non-photochemical quenching is very important in protecting PSII from damage to the photosynthetic mechanism in winter (Verhoeven *et al*., 1999; Demmig-Adams and Adams, 2000; Gilmore and Ball, 2000; Verhoeven, 2014). The radiationless dissipation of excess light occurring within the PSII reaction center was enhanced and could also contribute to the PSII photoprotection during winter (Gilmore and Ball, 2000; Ivanov *et al*., 2002; Gilmore *et al*., 2003; Yokono *et al*., 2008). It has been reported that accumulation of PsbS and/or Elip-like proteins during winter can also play an important role in photoprotection (Savitch *et al*., 2002; Ebbert *et al*., 2005; Verhoeven *et al*., 2009; Zarter *et al*., 2010; Míguez *et al*., 2017).

In addition to non-photochemical quenching processes, PSI-dependent cyclic electron flow (CEF) has been also suggested to play a significant role in preventing the photosynthetic apparatus from photodamage (Takahashi *et al*., 2009; Ivanov *et al*., 2012) and PSI-dependent CEF was reported to be enhanced during winter (Manuel *et al*., 1999; Ivanov *et al*., 2001).

The overwintering plants also reduced the chlorophyll concentration but increased carotenoids or anthocyanins concentrations during winter to lower the extent of photoinhibitory damage (Ottander *et al*., 1995; Matsubara *et al*., 2002; Robakowski *et al*., 2005; Maslova *et al*., 2009; Hughes, 2011; Wong and Gamon, 2015).

Although the photosynthetic performance of native overwintering species during winter has been extensively studied (Adams *et al*., 2001; 2004; Öquist and Huner, 2003; Verhoeven *et al*., 2014; Míguez *et al*., 2015), little is known about the photosynthetic adaptability during winter in introduced overwintering species. It has been reported that the photoinhibitory response and adaptation strategy during winter are different between woody and herbaceous plants (Verhoeven *et al*., 1999; Savitch *et al*., 2002; Margesin *et al*., 2007; Míguez *et al*., 2017). For that reason, both woody and herbaceous plants were used in this research. The common native overwintering species, lacebark pine (*Pinusbungeana*) and winter wheat (*Triticumaestivum*), a woody and herbaceous plants, respectively, were compared to bamboo (*Phyllostachysglauca*) and lilyturf (*Ophiopogonjaponicus*), a woody and herbaceous plants, respectively, introduced from south of China.

Here, the photosynthetic performance and the adaptive response during winter in two native overwintering species and two introduced overwintering species were compared by analyzing the photosynthetic gas exchange, chlorophyll a fluorescence, 820-nm light reflection and the relative abundance of photosynthesis related protein.

## Materials and Methods

### Plant materials

Four overwintering species including two woody species and two herbaceous species were used in this study. The lacebark pine (*Pinusbungeana* Zucc. ex Endl.) and winter wheat (*Triticumaestivum* L.) are woody and herbaceous species, respectively, are common native overwintering plants in Tai’an, where this experiment was performed. The bamboo *(Phyllostachysglauca* McClure) and lilyturf (*Ophiopogonjaponicus* (Thunb) Ker-Gawl) are woody and herbaceous plants, respectively, which were introduced from the south of China and are very popular and widely used as landscape greenery in the north of China.

All plants were grown in the south campus of Shandong Agricultural University, Tai’an City, Shandong Province, China (N36°09′49.78″, E117°09′4.72″). Current-year leaves (needles) from exposed branches of 30-year-old pine tree and 5-year-old bamboo, as well as leaves of current-year lilyturf and winter wheat were collected for analysis at 9:30 am from October 2017 to March 2018. Dark-adapted (30 min) leaves were used for physiological measurements (Chl a fluorescence and 820-nm light reflectance). Additional leaves were rapidly weighed, frozen in liquid N2 and kept at −80°C for further analyses.

Seasonal variations of temperature, light intensity and daily illumination duration were obtained from a weather station located close to the study site (Fig. 1).

**Fig. 1.**
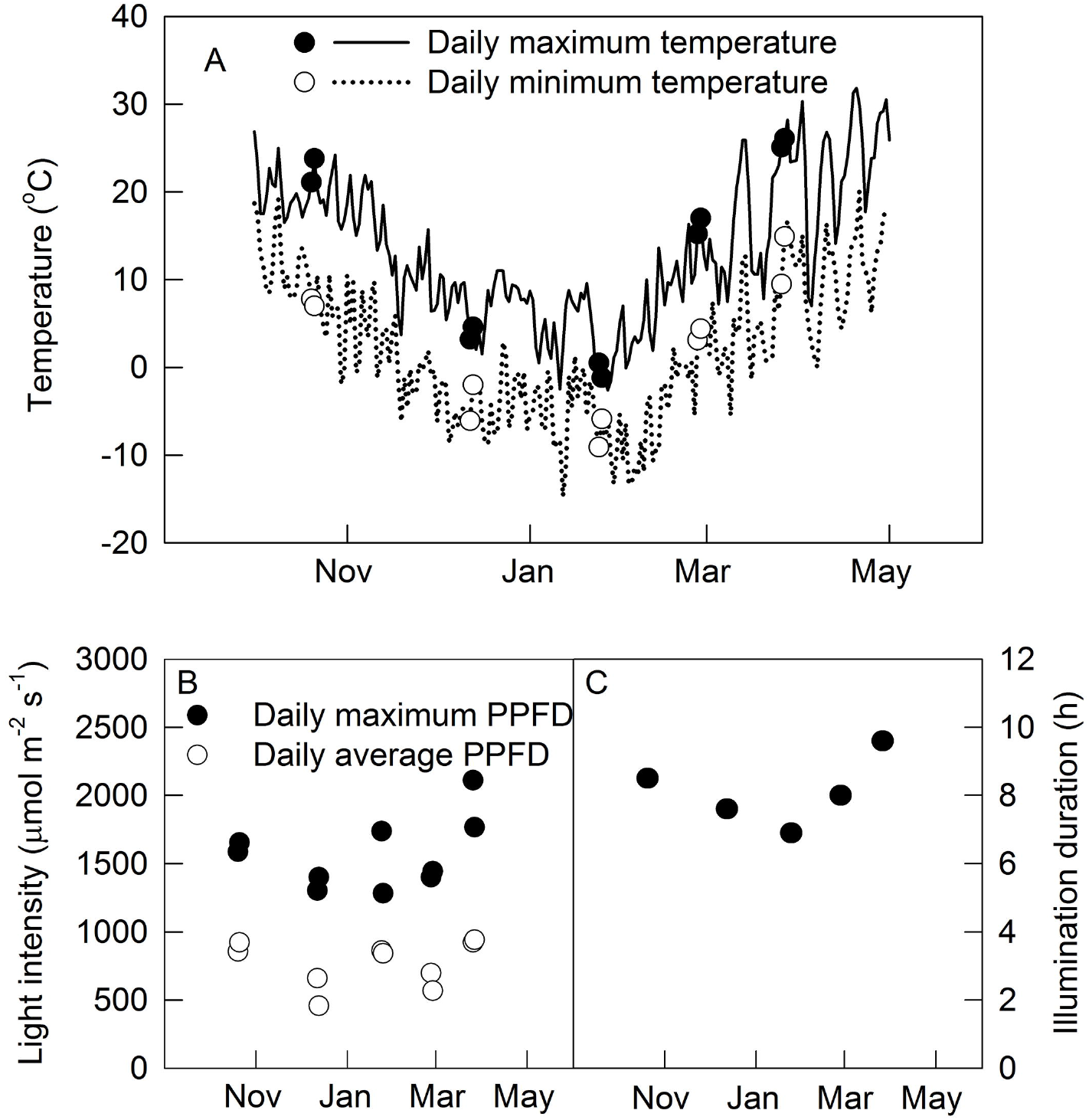
Seasonal variations of temperature, light intensity and illumination duration. Daily maximum (A, solid line) and minimum temperature (A, dashed line) from October 2017 to May 2018. The maximum (A, filled circles) or minimum temperature (A, open circles), the daily maximum light intensity (PPFD; B, filled circles), average light intensity (B, open circles) and illumination duration (C) at the day of the experiment.

### Determination of photosynthetic pigments

Frozen leaf samples were ground to a powder in liquid N2 and pigments were extracted by 80% (v/v) aqueous acetone. After centrifugation at 10000 x g for 5 min, the contents of Chl*a*, Chl*b* and total carotenoids in the supernatant were determined spectrophotometrically as described by Arnon (1949) and Porra *et al*. (1989).

### Photosynthetic gas exchange measurements

The net photosynthetic rates (Pn) in attached leaves were measured using a CIRAS-3 portable photosynthesis system (PP Systems International, Inc., Amesbury, MAUSA). The gas exchange measurements were performed from 10:00 to 15:00. The light intensity (1200 μmol m^−2^ s^−1^, 90% red light plus 10% blue light), CO_2_ concentration (400 μmol mol^−1^) and relative humidity (about 60%) were controlled by the automatic control device of the CIRAS-3 portable photosynthetic system.

### Measurements of the chlorophyll a fluorescence and 820-nm light reflection

The Chl a fluorescence and the 820-nm light reflection measurements on dark-adapted (30 min) leaves and needles were performed using an integral Multifunctional Plant Efficiency Analyser (M-PEA; Hansatech, UK) under ambient CO_2_ and O_2_ concentrations, and temperatures corresponding to the natural conditions as described earlier (Gao *et al*., 2014; Zhang *et al*., 2016). The dark-adapted leaves were illuminated by a two seconds saturating red light pulse (5000 μmol photons m^−2^ s^−1^) to obtain the maximum quantum yield of PSII (Fv/Fm). In addition, in dark-adapted leaves, the original value of 820-nm signal (Io) was firstlyrecorded, after that a twenty seconds of 10% far red light was applied, followed by a saturating red light pulse (1s, 5000 μmol photons m^−2^ s^−1^) to obtain the complete P700 oxidation (Pm, 100% P700^+^). The light induced P700 transient from Po to Pm (P700^+^) was used as a measure for the relative content of the active reaction centers of PSI (Klughammer and Schreiber, 2008).

### SDS-PAGE and immunoblot analysis

To extract soluble protein and thylakoid membranewereprepared in accordance with previous methods (Zhang *et al*., 2016; 2017). The thylakoid proteins contain 5 μg chlorophyll or 8 μL soluble protein supernatant were solubilized and separated by SDS-PAGE on 15% (w/v) acrylamide gels. Immunoblotting was performed by electrophoretically transferring the proteins from SDS-PAGE gels to nitrocellulose membranes (Bio-Rad Laboratories, Hercules, CA) according to the instruction book. The nitrocellulose membranes were blocked with 5% (w/w) skimmed milk and then incubated for 2 h with primary antibodies raised against the large subunit of Rubisco (1:5000), reaction centre protein of PSI, PsaA (1:2000), the reaction centre PsbA protein of PSII, PsbA(1:2000), the light harvesting proteins of PSI, Lhca1(1:2000), the major light harvesting protein of PSII complex (LHCII), Lhcb1(1:2000) and PsbS protein (1:2000). Subsequently, the membranes were incubated with horseradish peroxidase-conjugated anti-rabbit IgG antibody (Solarbio, Beijing, China) for 2 h. Immunoreaction of specific polypeptides was detected by using Supersignal West Pico substrate (Termo Fisher Scientifc, Shanghai, China) chemiluminescence detection kit and the immunoblots were visualized by using a Tanon-5500 cooled CCD camera (Tanon, Shanghai, China). The primary antibodies of Rubisco and PsbA were purchased from GenScript Co. Ltd. (Nanjing, China). The primary Lhcb1, PsbS, PsaA, and Lhca1 antibodies were purchased from AgriSera AB (Vanas, Sweden). Densitometric scanning and quantitative analysis of each replicate immunoblot was performed with a ImageJ 1.48v densitometry software (Wayne Rosband, National Institute of Health, USA, http://rsb.info.nih.gov.ij).

### Statistical analysis

All data points represent mean values ± SD calculated from 3-20 independent measurements. Tukey-Kramer’s method was used to analyze differences between the treatments using SPSS 11.

## RESULTS

### Environmental factors

Typical winter season in Tai’an, the investigation site, includes December, January and February. The average daily lowest temperatures were below 0°C during winter but higher than 0°C before December and after February. On the coldest day the measurements were performed (Jan. 24 2018) the daily lowest temperature was −9.1°C and the daily highest temperature was 0.5°C (Fig. 1). The illumination duration on Jan. 24 was also the shortest. The daily maximum light intensity was independent of the season.

### Pigment content

Chlorophylls are essential for the light absorption or light harvesting within the LHC polypeptides of PSII and PSI, excitation, energy transfer to the reaction centers of PSII and PSI and transformation of light energy to charge separated states within the reaction centers of PSII and PSI. Carotenoids not only act as supplementary light-harvesting pigments, but also play an important role in protecting the photosynthetic apparatus from the harmful effects of reactive oxygen species (ROS), especially singlet oxygen (Sandmann *et al*., 2014). The total chlorophyll (Chl a + Chl b) content of the native species did not exhibit seasonal variability in winter wheat Fig. 2A), but was decreased by 20-25% in lacebark pine during the winter season (Fig. 2B). In contrast, the introduced species exhibited a distinct seasonal pattern demonstrating almost 20% and 45% decrease of total chlorophyll content in lilyturf and bamboo leaves respectively, during the winter followed by its gradual and complete recovery during the spring months of March and April (Fig. 2A, B). The ratio of Chl a to Chl b (Chl a / Chl b; Fig. 2C, D) kept stable during winter in all four species used in this study.

**Fig. 2.**
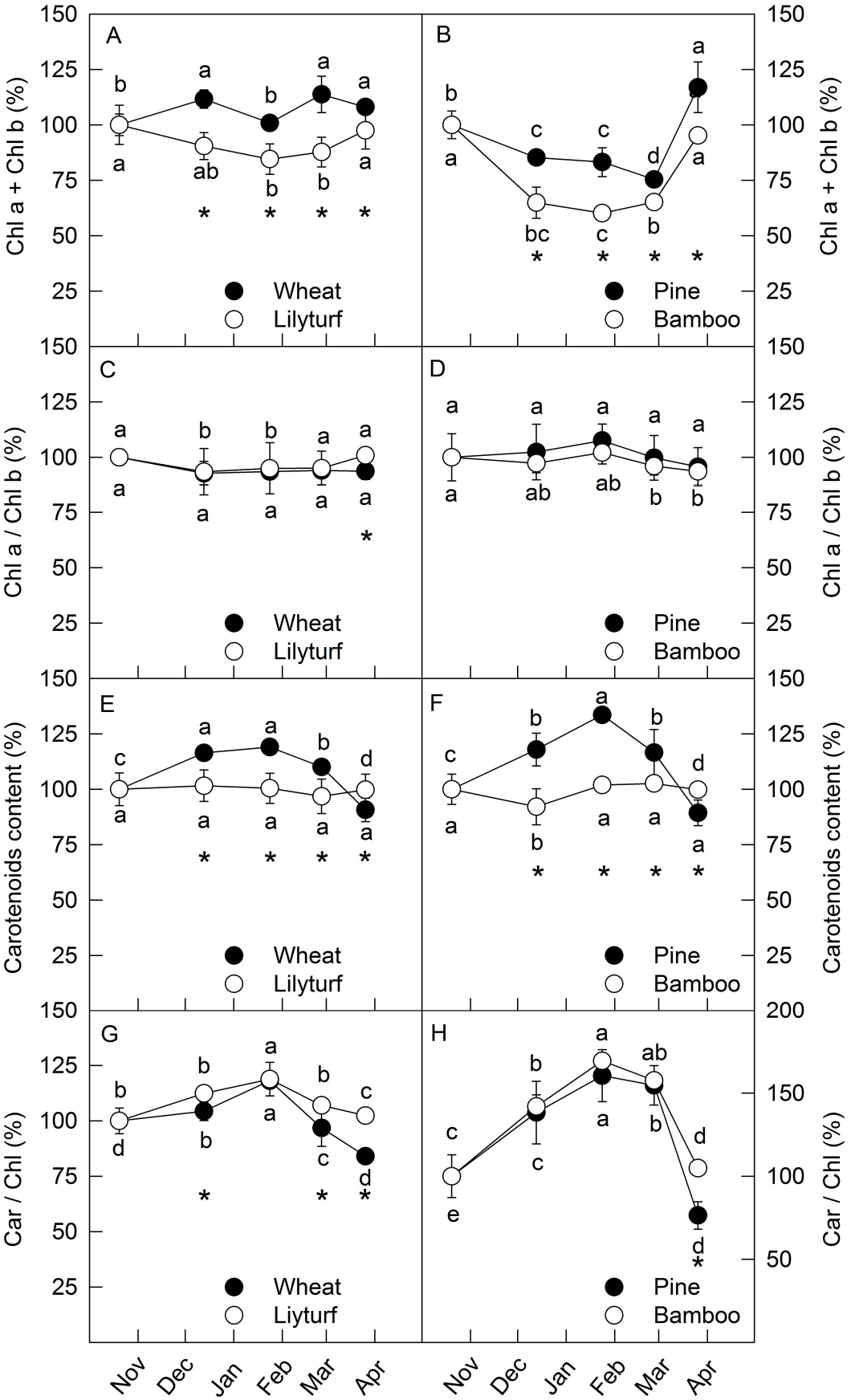
Seasonal variations of pigment in native overwintering species and introduced overwintering species. Total chlorophyll content (Chl*a* + Chl*b*; A, B), the ratio of Chl*a* to Chl*b* (Chl*a* / Chl*b*; C, D), the carotenoids content (E, F) and the ratio of carotenoids to chlorophyll content (Car / Chl; G, H) in two native overwintering species, lacebark pine (*Pinus bungeana*) and winter wheat (*Triticum aestivum*), as well as two introduced overwintering species, bamboo (*Phyllostachys glauca*) and lilyturf (*Ophiopogon japonicus*). The results are presented as percentages from the values observed on Oct. 20 (100%). Actual experimental data are shown in Supplementary Figure S1. All data points represent mean values ± SD calculated from 5 independent measurements. Different letters indicate significant differences at P <0.05 between different time. The asterisk indicates significant differences at P <0.05 between different species.

The total carotenoids content of leaves/needles in both native species demonstrated clear seasonal response (Fig. 2C, D). The carotenoids content was significantly increased by 20% (winter wheat) and almost 40% (lacebark pine) during the winter, while no significant seasonal changes in carotenoids content of both introduced species were observed (Fig. 2C, D). The ratio of carotenoids to chlorophyll content (Car / Chl; Fig. 2G, H) increased significantly during winter in all four species, but the change of this ratio was similar in native and introduced species.

These results imply that the native species are able to maintain higher light absorption and photoprotection capacity during winter compared to the introduced species.

### Photosynthetic CO_2_ fixation

The net photosynthetic rates (Pn) of both pine and bamboo woody plants were completed inhibited during winter and sharply recovered during the spring (April) (Fig. 3B). In contrast with woody plants, the photosynthetic CO_2_ fixation is still active during winter in both herbaceous species, although the Pn was significantly lower in winter compared to the Pn values registered in autumn and spring (Fig. 3A). The Pn decreased by 80~84% during winter in lilyturf but only 33~41% in winter wheat (Fig. 3A).

**Fig. 3.**
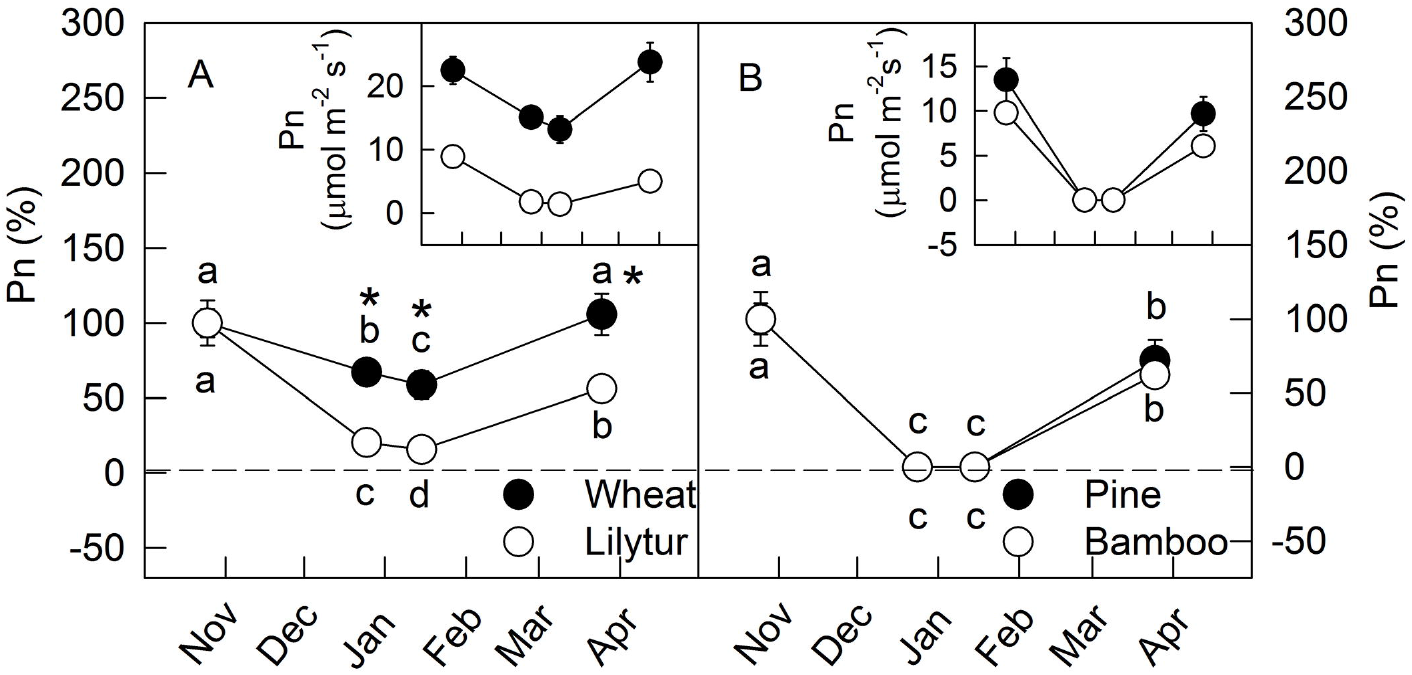
Seasonal variations of the net photosynthetic rates (Pn) in native overwintering species and introduced overwintering species. Two native overwintering species pine (*Pinus bungeana*) and winter wheat (*Triticum aestivum*; A), as well as two introduced overwintering species bamboo (*Phyllostachys glauca*) and lilyturf (*Ophiopogon japonicus*; B) were used. The results are presented as percentages from the values observed on Oct. 20 (100%). The original values of Pn were listed in insert. All data points represent mean values ± SD calculated from 5 independent measurements. Different letters indicate significant differences at P <0.05 between different time. The asterisk indicates significant differences at P <0.05 between different species.

### PSII and PSI photochemical activities

The maximum quantum yield of PSII measured as F_v_/F_m_ and the oxidation of P700 to P700^+^ (Pm) measured as light induced changes of 820-nm reflectance reflecting the primary photochemical activity of PSII and the relative content of active PSI reaction centers, respectively, were used to assess the seasonal changes of both PSII (F_v_/F_m_) and PSI (Pm).

In agreement with a numerous earlier reports (Ottander *et al*., 1995; Lundmark *et al*., 1998; Ivanov *et al*., 2001; Porcar-Castell *at al*., 2008; Verhoeven *et al*., 2009; Pieruschka *et al*., 2014) the photochemical efficiency of PSII (F_v_/F_m_) decreased significantly reaching F_v_/F_m_ values of about 0.3 during winter and recovered sharply in the spring in both woody species studied (Fig. 4B). It should be mentioned that the seasonal responses of F_v_/F_m_ were practically undistinguishable for the native (pine) and introduced (bamboo) species (Fig. 4B). Interestingly, the effects of F_v_/F_m_ to winter chilling were less pronounced in winter wheat and lilyturf compared to the woody species (Fig. 4A). In addition, while PSII photochemsitry of lilyturf decreased by 40%, winter wheat was only marginally affected (Fig. 4A).

**Fig. 4.**
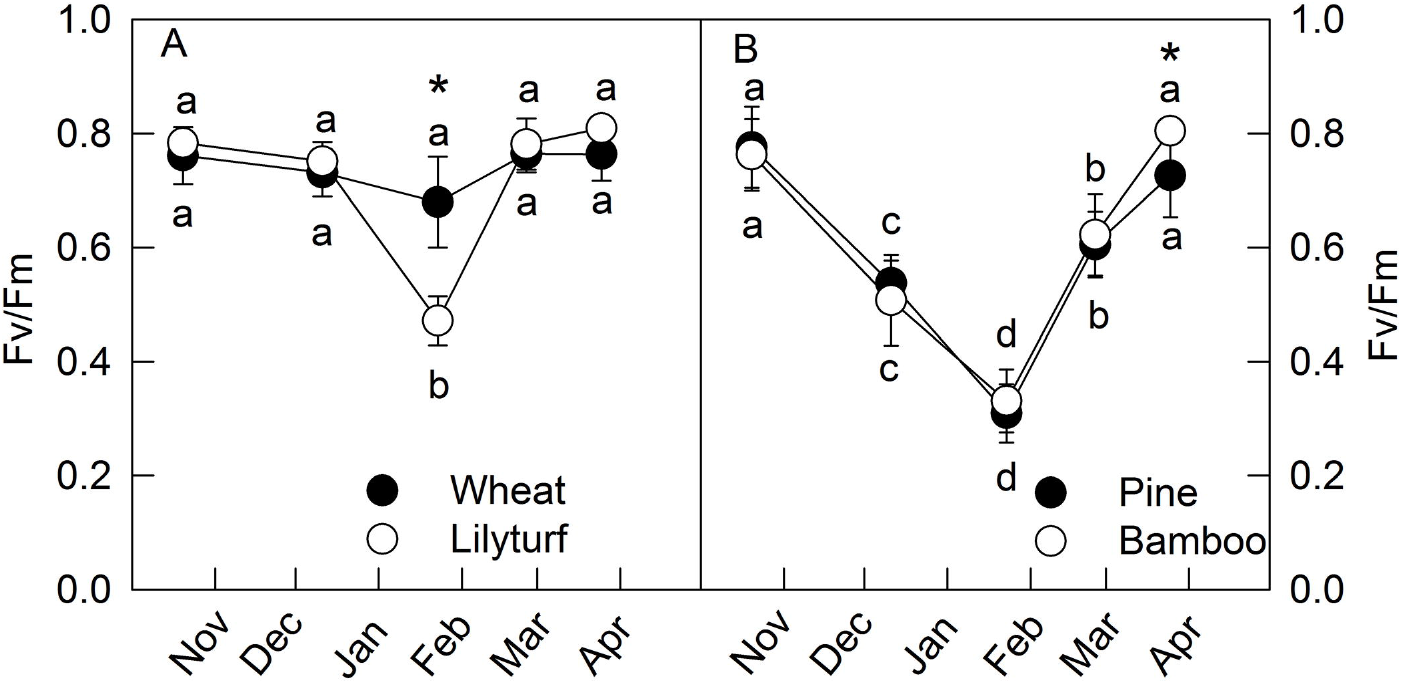
Seasonal variations of the maximum quantum yield of PSII (F_v_/F_m_) in native overwintering species and introduced overwintering species. Two native overwintering species pine (*Pinus bungeana*) and winter wheat (*Triticum aestivum*; A), as well as two introduced overwintering species bamboo (*Phyllostachys glauca*) and lilyturf (*Ophiopogon japonicus*; B) were used. All data points represent mean values ± SD calculated from 10 independent measurements. Different letters indicate significant differences at P <0.05 between different time. The asterisk indicates significant differences at P <0.05 between different species.

In addition to PSII photochemistry, seasonal changes of PSI photochemical performance were also followed by assessing the oxidation of P700 to P700^+^ (Pm). The extend of P700 photooxidation (Pm) exhibited minimal seasonal changes in winter wheat, while a significant decrease (35%) of Pm was registered inlilyturf during winter (Fig. 5A). In contrast to winter wheat and lilyturf, much stronger decrease (almost 70%) of Pm was registered in the woody species during the winter (Fig. 5B). It should be mentioned that while Pm values of pine recovered to 85% in March, the recovery of PSI photochemistry in bamboo was much slower and recovered to only 70% in April (Fig. 5B). These results clearly indicate that both PSI and PSII photochemical activities are more sensitive to winter photoinhibition in the introduced species (lilyturf and bamboo) compared to the native species (winter wheat and pine) studied.

**Fig. 5.**
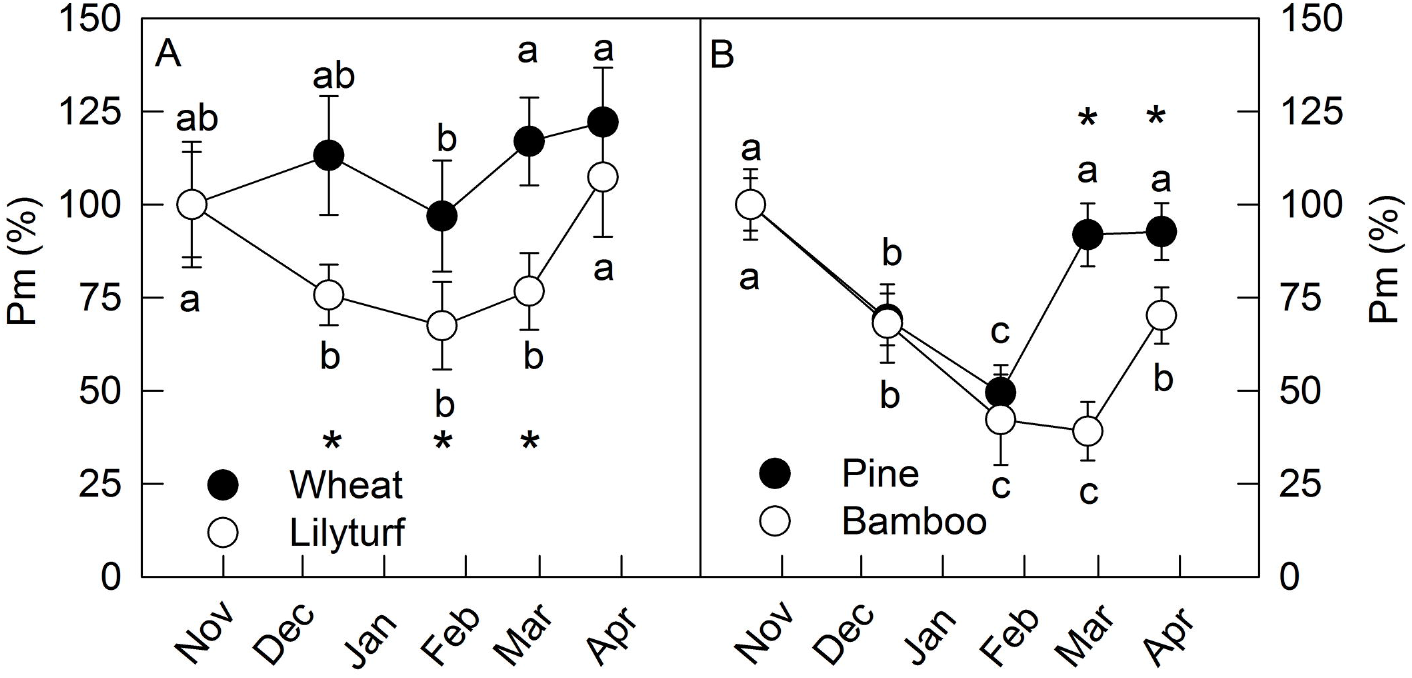
Seasonal variations of the relative content of the active PSI reaction centers (Pm) measured as P700^+^innative overwintering species and overwintering species. Two native overwintering species pine (*Pinus bungeana*) and winter wheat (*Triticum aestivum*; A), as well as two introduced overwintering species bamboo (*Phyllostachys glauca*) and lilyturf (*Ophiopogon japonicus*; B). The values are presented as percentages from the Pm (P700^+^) values observed on Oct. 20 (100%). Actual experimental data are shown in Supplementary Figure S2. All data points represent mean values ± SD calculated from 20 independent measurements. Different letters indicate significant differences at P <0.05 between different time. The asterisk indicates significant differences at P <0.05 between different species.

### Immunoblot analysis

The content of Rubisco, the key enzymes in photosynthetic carbon fixation was examined by immunoblot analysis. Representative immunoblots of the large subunit of Rubisco (RbcL) indicated that its relative abundance remained unchanged or even increased during winter in the two native species, i.e. pine and winter wheat, but was significantly reduced in the introduced species, lilyturf and bamboo (Fig.6A, D). The quantitative densitometric analysis also showed that the relative abundance of RbcL was marginally affected in winter wheat and pine, while in lilyturf and bamboo RbcL was decreased by 22.3% and 68.4%, respectively, during winter.

**Fig. 6.**
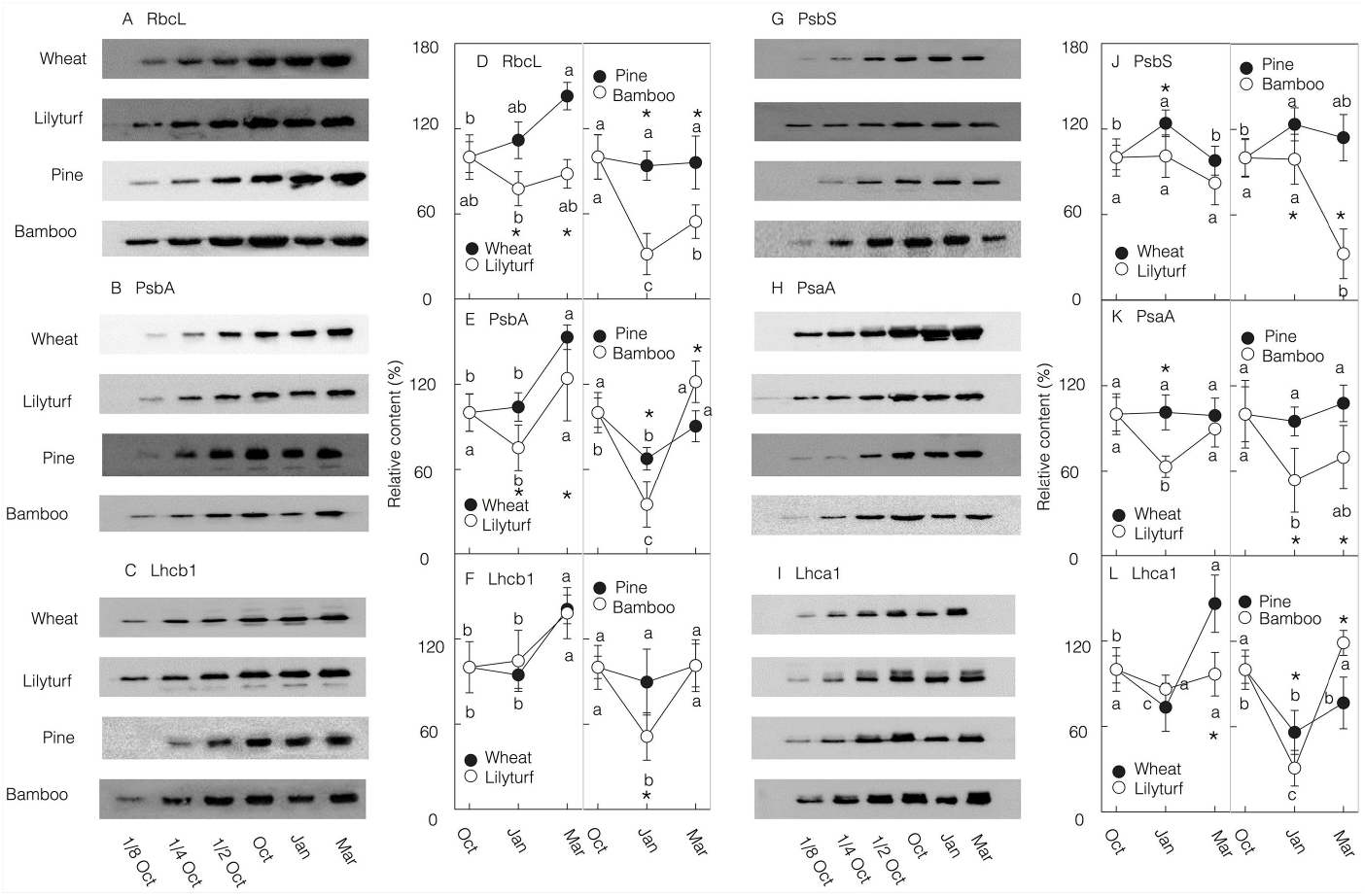
Seasonal variations of the photosynthesys related proteins innative overwintering species and introduced overwintering species. Two native overwintering species pine (*Pinus bungeana*) and winter wheat (*Triticum aestivum*; A), as well as two introduced overwintering species bamboo (*Phyllostachys glauca*) and lilyturf (*Ophiopogon japonicus*; B). The leaves were collected on Oct. 20 2017, Jan. 24 and Mar. 26 2018. The polypeptides were probed with specific antibodies raised against Rubisco large subunit (RbcL; A, D), PsbA (B, E), Lhcb1 (C, F), PsbS (G, J), PsaA (H, K) and Lhca1 (I, L). The typical immunoblots were presented in plots A-C and G-I, the data of quantitative densitometric analysis were presented in plots D-F and J-L. In plots d-f and J-L, all data points represent mean values ± SD calculated from three independent measurements. Different letters indicate significant differences at P <0.05 between different time. The asterisk indicates significant differences at P <0.05 between different species.

The relative abundance of PsbA (D1) (Fig. 6B, E) and PsaA (Fig. 6H, K) polypeptides, the core proteins of PSII and PSI reaction centers, respectively followed similar seasonal pattern. The relative amounts of PsbA (D1) and PsaA proteins, remained largely unaffected in native species during winter, while D1, was significantly reduced by 24.8% and 65.3% in lilyturf and bamboo, respectively (Fig. 6B, E). The amounts of PsaA were also decreased by 36.9% and 46.7% in lilyturf and bamboo, respectively, during winter season (Fig. 6H, K).

The representative immunoblots of the major constituents of the light harvesting Chla/b-protein complexes (LHC) associated with PSII (Lhcb1) and PSI (Lhca1) (Jansson, 1999) also demonstrated complex seasonal dynamics. The relative amount of Lhcb1 protein, the major subunit of LHCII, significantly decreased by 48.7% in bamboo during winter, compared to pine and both herbaceous species, where the amounts of Lhcb1 were marginally affected (Fig. 6C, F) In contrast, the content of LHCI subunit, Lhca1 protein, significantly decreased during the winter period in all species analyzed in this study (Fig. 6I, L).

PsbS protein is considered to play an essential role (Li *et al*., 2000) in developing the non-photochemical quenching (NPQ) of excess light energy, the most important photoprotective mechanism to PSII (Horton *et al*., 1996; Li *et al*., 2009; de Bianchi *et al*., 2010). The typical immunoblots presented in Fig. 6D and the quantitative densitometric analysis (Fig. 6J, G) clearly indicate that the relative content of PsbS protein increased by 24% and 23.5% in pine and winter wheat, respectively during winter, but remained unchanged in the introduced species (lilyturf and bamboo) (Fig. 6D).

## Discussion

This study provided evidences to prove that although some overwintering species have been successfully introduced to high latitudes from warmer areas, the photosynthetic adaptability to harsh winter is defective, and therefore cause the more serious photoinhibition in introduced species during winter, which may limit the growth and survival of introduced species at high latitudes.

### The lower photosynthetic CO_2_ fixation capacity in introduced species than native species during winter

This study showed that, the herbaceous species retain considerable photosynthetic CO_2_ fixation capacity but the Pn was approximately zero in woody species during winter (Fig. 3). Previous indoor research has reported that after an artificial 5°C cold acclimation, the Pn significantly decreased in lodgepole pine but was almost completely maintained in winter wheat (Savitch *et al*., 2002). The authors presumed that the photosynthetic adaptation strategies in winter are different between herbaceous and woody plants (Savitch *et al*., 2002). These earlier results generated in chamber experiment are confirmed in field experiments presented in this study. We also observed that Rubisco (RbcL) content was significantly reduced in both introduced species but remained almost unaffected in native species, regardless of herbaceous and woody species (Fig. 6A). The Rubisco possesses low catalytic efficiency and always restrict photosynthetic CO_2_ fixation (Raines, 2003). Therefore the lower amount of Rubisco in introduced herbaceous species may contribute to the more serious decrease of Pn during winter compared with native herbaceous species. Moreover, although CO_2_ fixation was completely inhibited during winter in woody species, the maintainence of relatively high amounts of Rubisco is still beneficial. Studies have shown that the synthesis of Rubisco requires nitrogen, and Rubisco accounts for up to 30% of total leaf nitrogen (Makino *et al*., 2003; Makino, 2011). To recover the CO_2_ fixation capacity during the next spring, the degraded Rubisco need to be re-synthesized in woody introduced species. Therefore, although the reduced Rubisco content in woody introduced species did not influence its winter CO_2_ fixation, it would delay the rapid recovery of photosynthesis during the next spring.

The degradation of Rubisco was also observed in cucumber and bean leaves during chilling-light condition, which was attributed to generation of excess reactive oxygen species (ROS) (Nakano *et al*., 2006; 2010). The ROS also took part in the Rubisco degradation under continuous mist or rain treatment under illumination (Ishibashi *et al*., 1996; Ishibashi and Terashima, 1996; Hanba *et al*., 2004; Nakano *et al*., 2006) and during senescence (Nakano *et al*., 2006; Feller *et al*., 2008; Ono *et al*., 2013). This experimental results presented in this study clearly demonstrate that both the PSII and PSI are more susceptible to photoinhibitions in introduced species than in native species (Fig. 4, 5). Earlier studies suggested that over-accumulation of ROS is one of the major reasons causing both PSI and PSII photoinhibiton (Choi *et al*., 2002; Sonoike, 2011; Zhang *et al*., 2014). So, we speculate that the more ROS caused by clod temperature and high light of winter in introduced species contributes to the degradation of Rubisco.

### The higher PSII photodamage and weaker PSII photoprotection in introduced species than native species during winter

The Fv/Fm decreased more serious in introduced herbaceous species than in native herbaceous species, but the decrease of Fv/Fm was almost the same in two woody species (Fig. 4). It should be noticed that sole analyzing the change of Fv/Fm is not enough to reflect the damage degree of PSII, which is due to both the PSII damage and photoprotection can decrease the Fv/Fm (Adams *et al*., 1994; Maxwell and Johnson, 2000; Verhoeven, 2013). The essential of PSII damage is the net degradation of D1 protein, the core protein of PSII reaction center (Vass, 2012; Yoshioka-Nishimura and Yamamoto, 2014). The capacity of NPQ is depend on the amount of PsbS protein and carotenoids, especially xanthophyll and lutein cycle pigments (Li *et al*., 2000; de Bianchi *et al*., 2010; Verhoeven, 2014). Therefore, we next analyzed the amount of D1 and PsbS protein. The D1 protein kept constant or slightly decreased and the PsbS protein content increased in native species, in contrast, the introduced species’s D1 protein decreased more obviously but the PsbS content was unchanged during winter. In addition, the carotenoids content increase in winter only in native species rather than in introduced species (Fig. 2). Above results proved that the reason caused the decrease of Fv/Fm was different between native and introduced species: the NPQ, one photoprotection mechanism, in native species but photodamage in introduced species.

Different from introduced species, the native species retained all chlorophyll (Fig. 2) and light-harvesting complex protein of PSII (Lhcb1; Fig. 6). Similar with Rubisco, the chlorophyll and light-harvesting complex also contained massive nitrogen (Makino *et al*., 2003) and therefore hard to re-synthesized during next spring. The reservation of chlorophyll and LHCII in native species will be beneficial for the rapid recovery of photosynthesis during the next spring. The reservation of chlorophyll and LHCII also indicates that the light absorbed by PSII during winter were higher in native species. Although the absorbed light was higher, the photodamage of PSII is alleviated in native species, which may be contributed by two mechanisms: (1) faster dissipation of light energy through NPQ and (2) more efficient utilization of light energy through CO_2_ fixation.

### Higher PSI photoinhibition in introduced species than native species during winter

During winter, the Pm decreased more obviously in introduced species than in native species (Fig. 5). Immunoblot analysis showed that the content of PsaA protein was unchanged during winter in native species, but decreased significantly in introduced species (Fig. 6). It was indicated that the PSI photoinhibition during winter was more serious in introduced species.

Although the Pm significantly decreased in pine, its PsaA content was maintained during winter (Fig. 4, 6). Previous studies have shown that in cucumber leaves exposed to chilling-light condition, the PSI photoinhibition occurred firstly in PSI receptors, the ferredoxin or iron-sulfur cluster (Sonoike *et al*., 1995b; Tjus *et al*., 1999; Teicher *et al*., 2000; Sonoike, 2006), the degradation of core protein of PSI reaction center occurred as the damage degenerated (Sonoike and Terashima, 1994; Ivanov *et al*., 1998; Tjus *et al*., 1999; Zhang *et al*., 2016). Therefore, we speculate that only ferredoxin or iron-sulfur cluster rather than PSI reaction center was damaged in pine during winter.

Active PSI is required for the cyclic electron flow around PSI (CEF). The CEF can generate a transmembrane proton gradient to activate NPQ and produce ATP that can be consumed in CO_2_ fixation (Yamori and Shikanai, 2016). In addition, the active PSI can also directly dissipate light energy in the form of P700^+^ (Kim *et al*., 2001; Ort, 2001; Bukhov *et al*., 2004; Suorsa *et al*., 2012), which may also function as an effective protective mechanism of the photosynthetic electron transport chain.

Maintaining the stability of the PSI reaction center complex is even more important than PSII, which is due to the recovery of PSI is much slower than PSII after photoinhibition. Studies have reported that re-synthesis of PSI reaction center protein is much slower than D1 protein after chilling-light induced photoinhibition (Kudoh and Sonoike, 2002; Zhang and Scheller, 2004). The Pm recovered much slower than the F_v_/F_m_ did, and the recovery of Pm is more sensitive to high light than F_v_/F_m_ (Zhang and Scheller, 2004; Zhang *et al*., 2011). During the next spring, the damaged PSI that can not recover quickly would limit the activity of the whole photosynthetic apparatus. Therefore, the inactivation of PSI and degradation of PSI reaction center protein will delay the recovery of photosynthesis during the next spring in introduced species.

This research also showed that the content of Lhca1 protein decreased in all four species during winter (Fig. 6). It was reported that under PSI photoinhibition treatment, the LHCI degraded earlier than PSI reaction center protein. And it was suggested that the degradation of LHCI is helpful for protection of PSI (Alboresi *et al*., 2009). In other words, the LHCI proteins act as fuses when other photoprotection mechanisms become insufficient (Alboresi *et al*., 2009). Our experimental results also imply that to protect the “precious” PSI reaction center during winter, both native and introduced species shared the same adaptive strategy: degrading LHCI that was “cheaper” and reducing the excitation energy to PSI reaction center.

In conclusion, the photosynthetic adaptability during winter in overwintering species introduced to higher latitudes are scarce and this study, for the first time, compared the photosynthetic adaptability during harsh winter between native and introduced overwintering species, including woody and herbaceous species. This study showed that the lower capacity for photosynthetic CO_2_ fixation and the more serious photoinhibition will endanger the survival of introduced overwintering species during winter; the degradation of photosynthetic related proteins will delay the recovery of photosynthesis during the next spring and therefore suppress the growth of introduced overwintering species.

## Supplementary information

**Supplementary Figure S1** Seasonal variations of pigment in native overwintering species and introduced overwintering species.

**Supplementary Figure S2** Seasonal variations of the relative content of active P700 in native evergreen species and introduced evergreen species.

## Acknowledgments

This study was supported by the National Natural Science Foundation of China project (31871544, 31701966, 31771691), Shandong Provincial Natural Science Foundation, China (ZR2017QC001) and Shandong Province Key Technology Innovation Project (2014GJJS0201-1).

